# Optimization of conidial production in the thermally dimorphic fungal pathogen *Histoplasma*

**DOI:** 10.64898/2026.08.20.745944

**Authors:** Bevin C. English, Murat C. Kalem, Mark Voorhies, Anita Sil

## Abstract

Sporulation is an integral process in the lifecycle of many fungal pathogens, including *Histoplasma*, a primary human pathogen that causes respiratory infections. *Histoplasma* conidia, or asexual spores, are the primary infectious particle but very little is known about them, in part due to the need for Biosafety Level 3 containment and inconsistency in generating viable conidia under laboratory conditions. Here, we identify media that consistently promote *Histoplasma* conidiation, yielding both micro- and macroconidia, and conditions that promote high levels of germination. We show that conidiation media and duration affect the proportion of macroconidia produced, and we demonstrate that *Histoplasma* strains vary in their response to these conidiation parameters. Finally, imaging studies of chitin, exposed chitin, and cell wall mannoproteins show that while micro- and macroconidia have similar cell wall compositions, strain type and conidiation media variation result in qualitative differences in staining. These optimized methods for *Histoplasma* conidial preparations will enable more detailed investigations into this understudied aspect of the biology of an important human fungal pathogen.

**Importance:** *Histoplasma* is a thermally dimorphic fungal pathogen that can infect a wide range of mammalian hosts. Unlike the majority of other fungal pathogens, such as *Candida* and *Aspergillus* species, *Histoplasma* can cause serious, sometimes fatal, disease in otherwise healthy, immunocompetent individuals. *Histoplasma* infections occur when asexual spores called conidia are inhaled by the mammalian host. Despite being the infectious particles, conidia are the least-studied morphologic form of the fungus, due in part to technical challenges in reliably producing conidia in the laboratory. In this work, we describe optimization of conidiation conidiations, including sporulation media and duration of conidiation, allowing us to interrogate the cell wall composition of conidia produced by different *Histoplasma* species.

## Introduction

*Histoplasma* is the etiological agent of histoplasmosis, a fungal disease that can affect both immunocompromised and immunocompetent individuals [1]. *Histoplasma* is one of the most common endemic mycoses in the Americas and is found throughout the world, and recent analyses suggest that its global range may be broader than previously thought [2]. *Histoplasma* is a thermally dimorphic pathogen: like other related thermal dimorphs, including *Blastomyces* spp., *Coccidioides* spp., and *Paracoccidioides* spp., *Histoplasma* grows in two distinct morphologies. At ambient temperatures in the environment, *Histoplasma* grows in the soil as a multicellular network of hyphae, which can produce asexual spores called conidia. When the soil is disrupted, these spores may become windborne, potentially settling into a new suitable environment and germinating into hyphal cells. Alternatively, these spores may be inhaled by a mammalian host. Mammalian body temperature is one key signal that triggers a morphogenic switch by the fungus, causing it to grow in the pathogenic yeast form [1, 3].

*Histoplasma* is unique among the thermal dimorphs in that it makes both microconidia and macroconidia. As their names suggest, these spores are differentiated by their size, with microconidia ranging from 2-5 μm in diameter and macroconidia ranging from 8-14 μm. Macroconidia are frequently tuberculate and are so distinctive that they are often helpful in identifying *Histoplasma* grown from patient samples [4]. Because particles under 5 μm can penetrate deeper in the lung than larger particles [5], microconidia are assumed to be the infectious spores [1, 4]; however, this has not been empirically tested. Indeed, it is unknown if micro- and macroconidia differ in any way other than size.

Though conidia are an important part of the *Histoplasma* life cycle, they are woefully understudied. In addition to differences between micro- and macroconidia, very little is known about what signaling pathways contribute to conidiation or germination. This lack of understanding is partly due to several technical reasons. Because the infectious spores pose an inhalation risk, they require Biosafety Level 3 (BSL-3) containment. Further, conidiation can be a long process, taking weeks to months. And finally, the difficulty in obtaining consistent conidial preparations with reliable germination rates has been noted for decades [6–8]. Thus, we set out to rigorously test *Histoplasma* conidiation parameters, with the goal of identifying optimal conditions to consistently produce large amounts of viable spores, ultimately enabling us to perform initial qualitative comparisons of cell wall compositions of micro- and macroconidia from different *Histoplasma* strains.

## Results

### 2x GYE, Malt, and V8 media induce *Histoplasma* conidiation

We selected 8 types of agar plates to evaluate as *Histoplasma* conidiation media, all of which are detailed in supplemental file 1: 2x glucose yeast extract (2x GYE), chicken soil (CS), modified cysteine-glutamine (Cys-Gln), malt extract, Sabouraud (SAB), soil, modified uBird, and V8. 2x GYE is routinely used for arthroconidia production by *Coccidioides* [9], a thermally dimorphic fungus closely related to *Histoplasma.* We created CS media because previous studies have shown that manure from starlings [10] and other animals [11] can stimulate conidiation in *Histoplasma*. We have previously observed that modified Cys-Gln media [12] supports *Histoplasma* conidiation [13]. Malt extract, Sabouraud, and soil agars are commonly used for culturing fungal species and have been previously used for *Histoplasma* sporulation [7, 8, 14]. Modified uBird is derived from Bird media, which was developed as an alternative media for culturing *Neurospora crassa* [15]. V8 agar is commonly used to culture fungi and is widely used for *Cryptococcus* sexual spore production [16]. For consistency, all plates were aged for two weeks at room temperature while protected from light, as described for V8 agar [16].

We inoculated the different media with yeast from the *Histoplasma* strain G217B and incubated them at 25°C for 31 days. Unsurprisingly, we observed varying macroscopic morphologies, including thick, felt-like hyphal mats on 2x GYE and SAB, thinner hyphal mats on soil and Cys-Gln, and dry, wispy growth on V8 (Fig S1). The different media also resulted in very different conidial preparations (Fig 1). Conidial suspensions from 2x GYE, malt, SAB, and V8 media were mixtures of micro- and macroconidia. CS, Cys-Gln, soil, and uBird resulted in preparations that contained conidia and other material, including elongated and ovoid cells and presumably dead phase-dark material (Fig. 1A); thus, these media were excluded from further investigation. SAB did not consistently yield enough spores to accurately quantify and was thus also excluded from further study (Fig 1B). *Histoplasma* grown on V8 and malt agar produced comparable conidial yields, which were significantly higher than those isolated from 2x GYE (Fig 1B). These conidial preparations also differed in their percentage of macroconidia, with 2x GYE and V8 media producing significantly more macroconidia than malt agar, which produced less than 5% macroconidia (Fig 1C).

**Figure 1.**
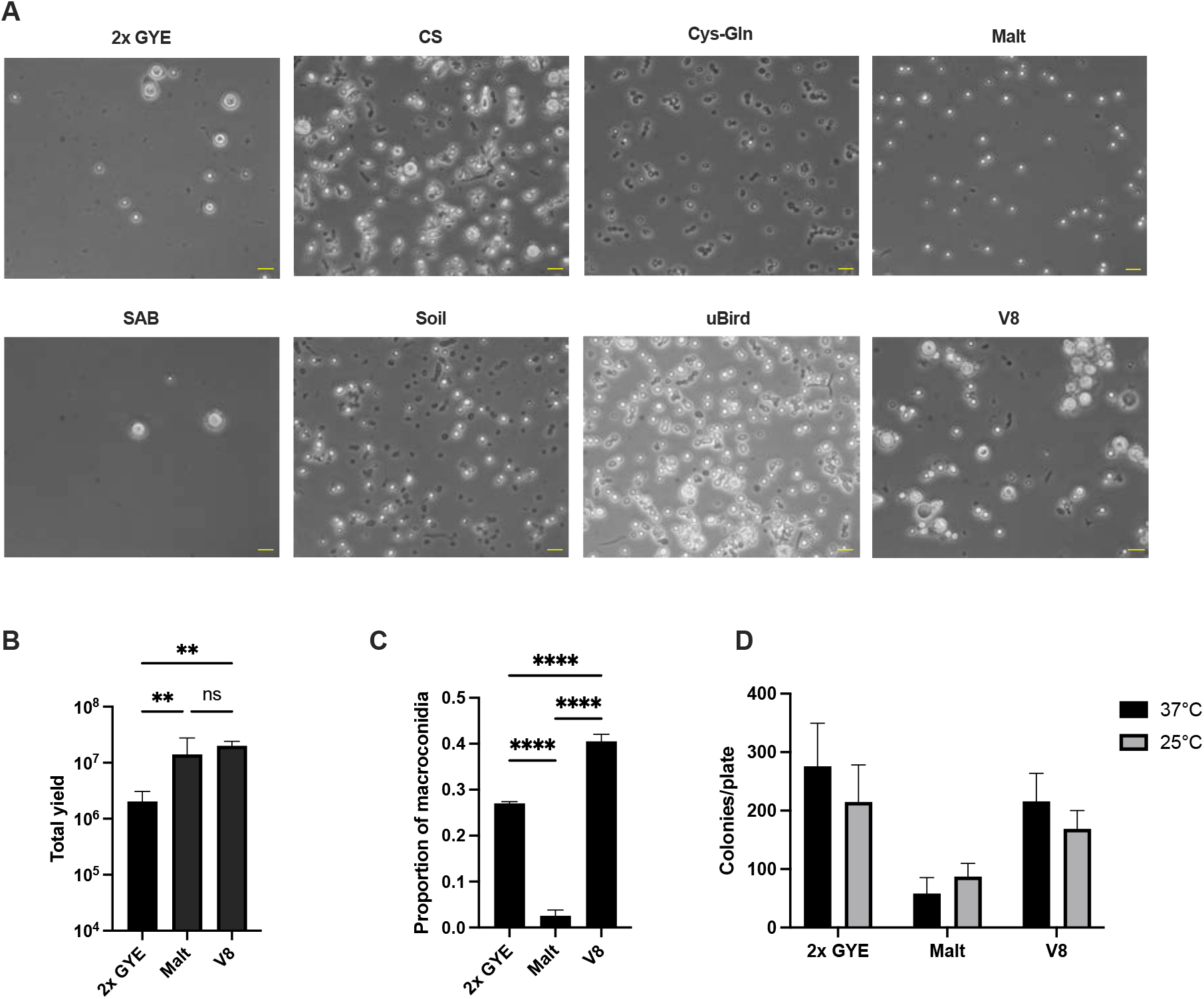
Comparison of conidiation media. *Histoplasma* yeast were inoculated on agar plates of the indicated media, and conidia were prepared after 4 weeks at 25°C. (A) Representative images of conidial preparations. Yellow scale bar, 10 μm. (B) Total conidia collected from the indicated media types. Only 2x GYE, malt, and V8 yielded consistent preparations of sufficient quantities for enumeration. **, p< 0.01, one-way ANOVA with Tukey’s multiple comparison’s test on log-transformed data. (C) Relative proportion of macroconidia prepared from the indicated media. ****, p< 0.0001, one-way ANOVA with Tukey’s multiple comparison’s test. (D) For each conidial preparation, 500 spores were plated on BHI agar. Colonies were enumerated after 2 weeks at 25°C or 37°C. Statistical analysis results are shown in Supplemental Table 1.

We noticed that the spores prepared from V8 plates formed large aggregates that were not disrupted by vortexing (Fig 1A). Because we do not understand the mechanism underlying the adhesion, we tested a variety of chemicals to aid in disaggregation: Tamol is a chemical dispersant used in *Coccidioides* spherulation media [17] and is necessary for efficient endospore dispersal [18]; mannose and EDTA have been shown to inhibit flocculation in *Saccharomyces cerevisiae* [19]; Tween-80 and Triton X-100 are common nonionic detergents, with Tween-80 generally considered a milder detergent. Only 0.5% Tamol and 0.1% Triton X-100 were able to disrupt the spore clumps (Fig S2A). Because 0.1% Triton X-100 was quite bubbly and thus posed an aerosol risk in the BSL-3, we opted to further examine the effects of Tamol. As expected, diluting spores in Tamol increased plating efficiency (Fig S1B), presumably by disaggregating the fungal cells, though we cannot rule out other effects at this time.

We assessed the viability of spores generated on different conidiation media. For spores generated on 2x GYE and V8, we observed high germination rates, with approximately 50% of spores of germinating at 37°C; we also observed a trend toward more germination at 37°C compared to 25°C, though this was not statistically significant. Spores from these media types also had higher germination rates than those generated on malt, which showed higher germination rates at 25°C than 37°C but had less than 20% germination (Fig 1D; Supplemental Table 1).

### Length of conidiation incubation has mild effects on yield and proportion of macroconidia

Length of time on conidiation plates has been shown to influence a range of spore characteristics [14, 20], including the presence of microconidia in *Neurospora crassa* [21]. Thus, we set out to determine how incubation time affected *Histoplasma* strain G217B conidiation on V8 plates. We were able to isolate conidia from *Histoplasma* grown for 2 to 8 weeks; however, we noticed that longer incubation periods led to more phase-dark debris in our preparations (Fig 2A). The yields obtained at 2 weeks post-inoculation were highly variable. For the subsequent time points, length of conidiation time correlated with yield, though the effect size was small (Fig 2B). Intriguingly, time of conidiation also correlated with percentage of macroconidia in the preparation, though again the effect size was relatively small (Fig 2C). Consistent with our previous observations (Fig 1D), more spores germinated frequently at 37°C compared to 25°C (Fig 2D). Unsurprisingly, germination rates varied between experiments (Fig S3), but there was no increase in germination rates after 4 weeks of conidiation (Fig 2D). Together, these data show that while there is a slight increase in yield and in macroconidial content after 4 weeks of conidiation, there is more debris and no increase in germination rates, leading us to conclude that 4 weeks of conidiation on V8 plates is optimal for *Histoplasma* strain G217B.

**Figure 2.**
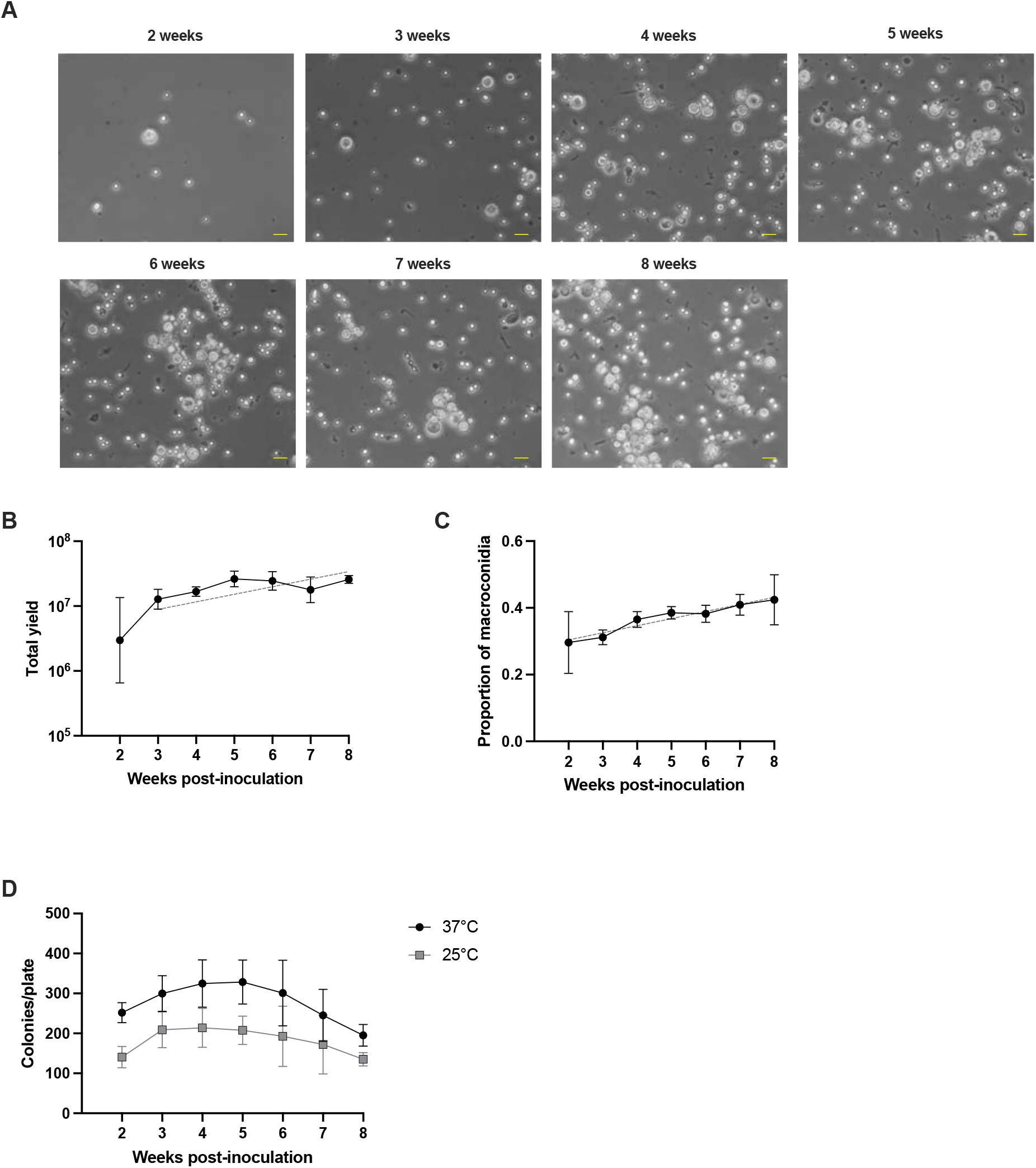
Conidiation time affects yield and composition but not germination. *Histoplasma* yeast were inoculated on V8 agar plates and incubated at 25°C for 2-8 weeks before conidia were prepared. (A) Representative images of conidial preparations. Yellow scale bar, 10 μm. (B) Total conidia collected at the indicated times. Simple linear regression of yields from weeks 3 to 8 is shown as a grey dashed line (slope = 0.1168; 95% confidence interval 0.04403 to 0.1896). (C) Relative proportion of macroconidia prepared after the indicated times. Simple linear regression shown as grey dashed line (slope = 0.02125, 95% confidence interval 0.01110 to 0.03141). (D) For each preparation, 500 spores were plated on BHI agar and colonies were enumerated after 2 weeks of growth at 25°C or 37°C.

### *Histoplasma* strains vary in conidiation characteristics

*Histoplasma* is found throughout the world and was recently divided into five genetically distinct lineages that differ in geographical regions of endemicity [22]. Previous studies have indicated that sporulation varies between *Histoplasma* isolates [8, 23], and we thus wondered how strains from different lineages would respond to our conidiation conditions. In addition to G217B, which was recently reclassified as *Histoplasma ohiense*, we examined G184AR, which belongs to the Panama lineage of *Histoplasma capsulatum*, and H88, which belongs to the Africa lineage *Histoplasma capsulatum* var. *dubosii* [22]. Unsurprisingly, these strains exhibited different macroscopic morphologies on both V8 and malt agar plates: compared to G217B, G184AR appeared whiter and fluffier on V8 but had noticeably fewer aerial hyphae on malt plates. H88 had white hyphal mats that were noticeably fluffier than the other strains on both media types (Fig S4). We were unable to isolate any spores from H88, even if the conidiation plates were incubated for 9 weeks; however, we were able to collect conidia after 4 and 6 weeks of growth on both media from G217B and G184AR (Fig 3A). Most preparations resulted in comparable yields, though the yield trended lower for G184AR grown on malt plates for 4 weeks. Consistent with our previous observations, we obtained more spores from V8 agar than malt agar, and we obtained more spores after 6 weeks of conidiation compared to 4 weeks (Fig 3B, Supplemental Table 2). Consistent with our previous results, G217B grown on V8 plates produced a large amount of macroconidia. However, G184AR produced very few macroconidia on V8 plates; in fact, the higher macroconidial yield from G184AR was observed from malt plates (Fig 3 A, C). Also consistent with our previous results, we saw higher germination rates at 37°C than at 25°C, and spores generated on V8 plates had higher germination rates than those from malt plates (Fig 3D; Supplemental Table 3).

**Figure 3.**
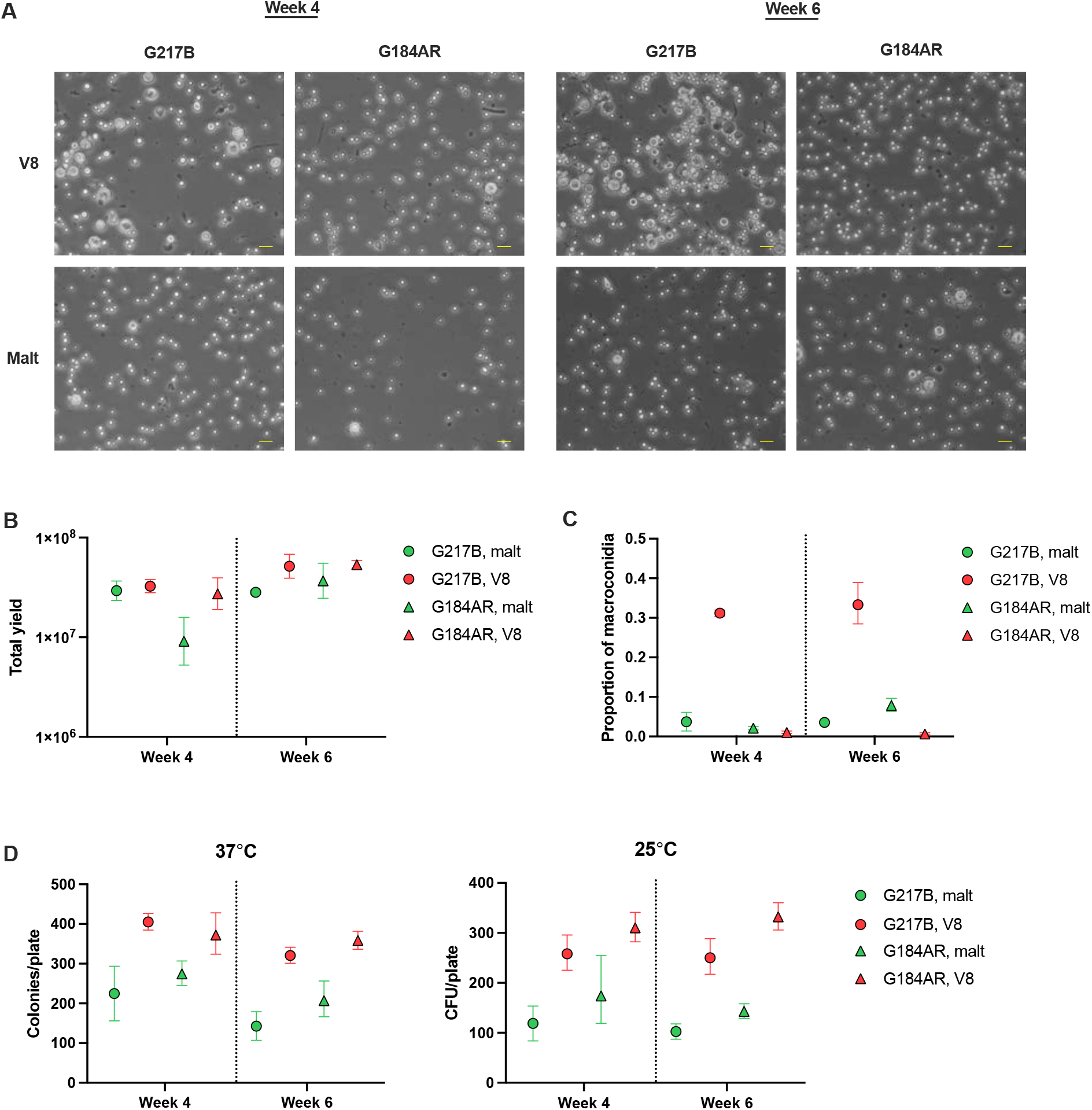
Conidial preparations from *Histoplasma* strains vary in yield, composition, and germination. *Histoplasma* yeast from strains G217B and G184AR were inoculated on V8 and malt agar plates. After 4 and 6 weeks of incubation at 25°C, conidia were prepared. (A) Representative images of conidial preparations. Yellow scale bar, 10 μm. (B) Total conidia collected in the indicated preparations. Statistical analysis results are shown in Supplemental Table 2. (C) Relative proportion of macroconidia in the indicated preparations. Proportions of macroconidia on V8 vs malt were compared independently for each strain, time, and batch using the prop.test function in R. All eight tests were significant with p < 2e-16. (D) For each preparation, 500 spores were plated on BHI agar, and colonies were enumerated after 2 weeks of growth at 25°C or 37°C. Statistical analysis results are shown in Supplemental Table 3.

### Conidial cell wall composition differs by media and strain

The cell wall is critical for structural integrity, resistance to harsh environmental conditions, and host-pathogen interactions [24–27]. Therefore, we reasoned that cataloging differences in conidial cell wall composition across strains and media types would be a strategic first step in characterizing conidial cell biology. The conidial cell wall composition was examined using fluorescent probes. These include calcofluor white, wheat germ agglutinin, and concanavalin A to determine the presence and distribution of total chitin, exposed chitin, and mannoproteins, respectively. Our analysis focused on G217B and G184AR conidia prepared from malt and V8 plates after 4 weeks post-inoculation (Fig 4). Both micro- and macroconidia cell walls had an inner layer of chitin decorated with exposed chitin and mannoproteins. We observed a punctate staining pattern for mannoproteins in both G217B (Fig 4A) and G184AR (Fig 4B) micro- and macroconidia. G184AR conidia also had robust, punctate exposed chitin staining. The punctate pattern in macroconidia mostly colocalized to tuberculate protrusions. The cell wall of G217B conidia had qualitatively less exposed chitin and mannoproteins compared to G184AR in malt. Comparison of the exposed chitin between malt and V8 media revealed that conidia from both G217B and G184AR grown on malt had more exposed chitin, though it was more pronounced in G184AR. Importantly, we noticed an appreciable degree of heterogeneity in cell wall composition across conidia. For example, the G217B V8 conidial preparation had a macroconidium with a thick chitin layer and high mannoprotein staining (Fig 4A, yellow arrow) and another macroconidium had a thin layer of chitin with less mannoprotein staining (Fig 4A, orange arrow). Similar observations, including a high degree of heterogeneity, were seen with spores produced after 6 weeks of conidiation (Fig S5).

**Figure 4.**
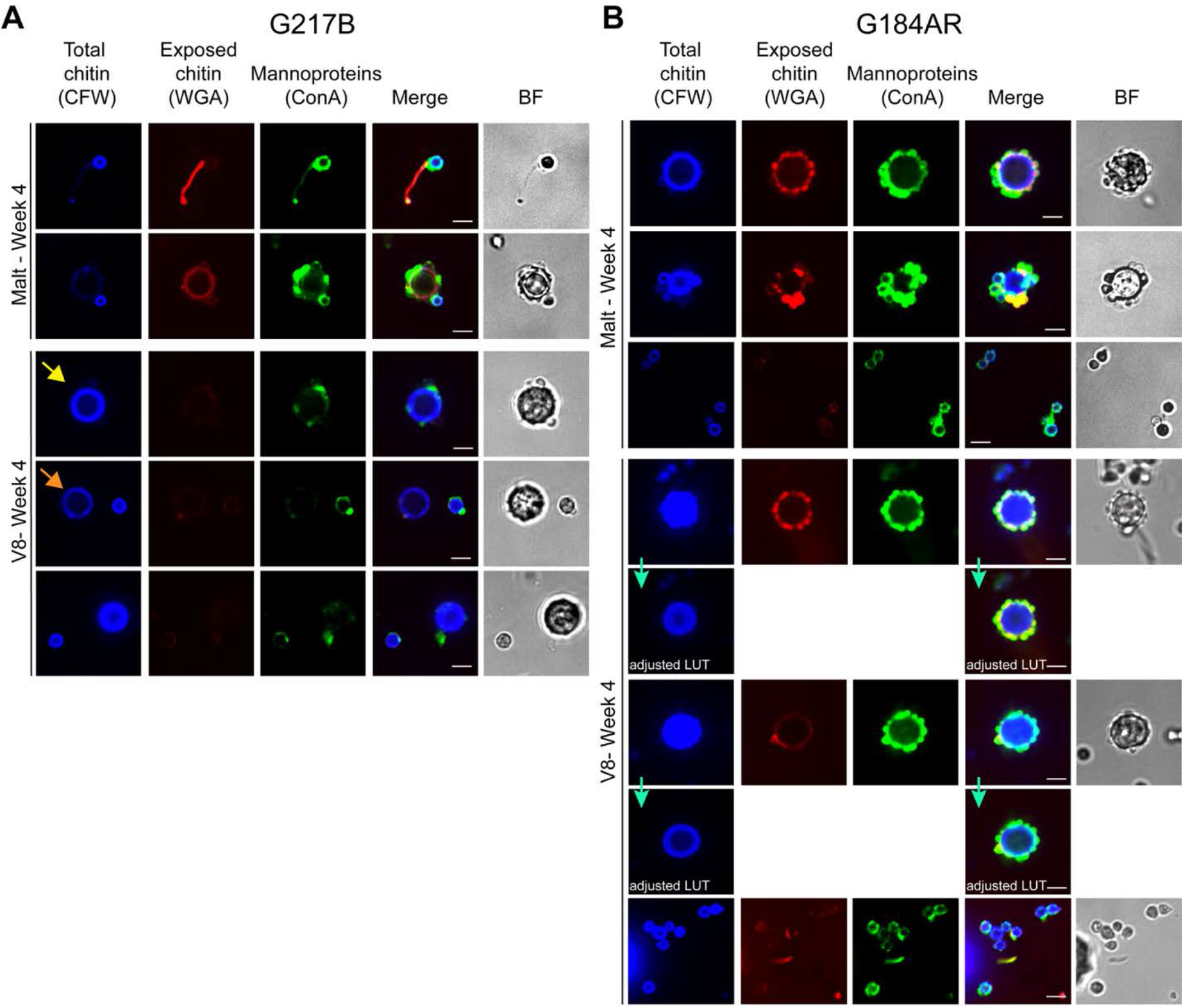
Conidial cell wall composition is influenced by *Histoplasma* strain and conidiation media. *Histoplasma* yeast from strains G217B and G184AR were inoculated on malt and V8 agar plates. After 4 weeks of incubation at 25°C, conidia were prepared. Conidia were fixed and stained with calcofluor white (CFW), wheat germ agglutinin conjugated to Alexa Fluor 594 (WGA), and concanavalin A conjugated to Alexa Fluor 488 (ConA) to detect cell wall chitin, exposed chitin, and mannoproteins, respectively. Representative images for (A) G217B and (B) G184AR conidia are shown. Images indicated by green arrows show LUT (look-up table) adjustments alongside unadjusted images to improve visualization (green arrows). Yellow and orange arrows highlight macroconidia with thick and thin chitin layers, respectively. BF, bright field. White scale bars, 5 µm.

## Discussion

Conidia are an important component of the *Histoplasma* lifecycle and are the infectious form; however, they are also the least studied morphological form, due in part to the difficulty in obtaining consistent conidial preparations with reproducible germination rates, which has been noted for decades. Thus, we set out to optimize conidiation conditions to increase reproducibility.

One of the most important parameters for conidiation is the growth medium. In this study, we identified 2x GYE, malt, and V8 agar media that reliably promote conidiation of two different *Histoplasma* species. For consistency, we aged all conidiation plates, as previously described for V8 agar [16]. In the past, *Histoplasma* conidiation plates have been sealed with Parafilm after inoculation [8, 14]. However, in addition to aging the plates, the other factor that led to more consistent conidial preparation was keeping the plates unwrapped during conidiation. We hypothesize that one factor leading to the reduction in variability is more uniform humidity as a result of both drying during aging and leaving them unwrapped after inoculation. Interestingly, it has been demonstrated that drying V8 agar plates in a biosafety cabinet does not fully recapitulate the effect of aging plates [16]; thus, other factors may be contributing to the improvement in consistency. Of course, this was not an exhaustive study of all possible conidiation media, and other media may also promote *Histoplasma* sporulation. However, as 2x GYE, malt, and V8 agar are all straightforward to produce, we recommend using aged, unwrapped agar plates of these three media for *Histoplasma* conidiation.

In addition to growth medium, incubation time is another important parameter for conidiation. We were able to isolate spores from *Histoplasma* strain G217B grown on V8 agar for as little as two weeks, though maximum yield was not achieved until 4 weeks of conidiation. However, for strain G184AR grown on malt plates, 4 weeks was not sufficient for maximal yields, as we obtained more spores after 6 weeks of growth. We also observed that incubation time correlated with the amount of phase-dark debris in preparations from G217B incubated on V8 agar. Longer conidiation time correlated with an increase of macroconidia for both G217B grown on V8 agar and G184AR grown on malt. We have previously shown that *Histoplasma* conidia, but not yeast cells, induce a type I interferon response in bone marrow-derived macrophages. Intriguingly, the magnitude of this response correlated with the duration of conidiation [14]. Thus, we believe that while future studies may need to vary conidiation incubation times depending on the *Histoplasma* strains, researchers should maintain consistent conidiation incubation periods whenever possible.

We were unable to isolate any spores from H88, an African strain of *Histoplasma*, even after incubation on V8 and malt plates for 9 weeks. It is unknown if anyone has ever successfully prepared spores from H88. We and others have previously failed to isolate conidia from other *Histoplasma* strains on different sporulation media [8, 23]. It is possible that some laboratory strains may have lost the ability to conidiate, or we simply have not identified universal conidiation conditions for highly diverged *Histoplasma* strains. As our knowledge of the signaling pathways that lead to sporulation grows, we may be able to distinguish between these two possibilities, but that is outside the scope of the current work.

When evaluating viability, we chose to germinate the spores on brain heart infusion (BHI) agar plates supplemented with sheep’s blood. We observed a much higher germination rate on BHI (Fig. 1-3) than we have previously observed on HMM plates [13], which are commonly used to grow *Histoplasma* yeast and hyphae. BHI plates may mimic host conditions with regard to nutrient availability, and these host-like conditions may contribute to increased germination at 37°C. Spores from different fungi, including the related thermally dimorphic fungus *Talaromyces marne>ei*, have differential germination capabilities that correspond to conditions during conidiation [28]. Intriguingly, we have previously reported that spores generated on Cys-Gln media are unable to germinate on that same media at 25°C but can at 37°C [13]. More work is needed to understand the factors that promote germination of *Histoplasma* conidia. Many previous studies have noted wildly variable germination rates [6, 7, 10, 29]; however, we were able to observe more consistent germination rates across experiments, and thus we recommend BHI plates supplemented with sheep’s blood for germination. As conidia may form aggregates, we also recommend using the dispersant Tamol to maintain a single cell suspension, increasing accuracy when assessing germination rates via CFU enumeration.

After exploring optimal conidiation conditions, we wanted to more closely examine the conidia we generated under those conditions. To this end, we examined the cell wall compositions of the different spores we isolated. We noted qualitative cell wall differences across media types and strains. We speculate that the cell wall differences across media types could be due to differences in nutrient availability and sugar metabolism [30]. *Histoplasma* strains are classified into two chemotypes based on yeast cell wall polysaccharide composition [31]. The observed qualitative differences in conidial cell wall across strains might be explained by the reported differences across chemotypes. While yeast cells of chemotype II strains (e.g., G184AR) contain α-(1,3)-glucan, chemotype I strains (e.g., G217B) do not [32]. Presence or absence of α-(1,3)-glucan likely alters the abundance of other cell wall components. Lastly, we observed a high degree of cell wall heterogeneity, suggesting that individuals within the population of spores could exhibit phenotypic variation.

The ultimate goal of our work optimizing *Histoplasma* conidiation conditions is to enable further study of this important feature of the *Histoplasma* lifecycle. Very little is known about the regulatory pathways that govern conidiation. We have previously shown that key transcription factors that regulate *Histoplasma* morphology are also involved in conidiation, including Stu1, Fbc1, Pac2 [33], Ryp1[34], Ryp2, and Ryp3 [35]. The histidine kinase Drk1 may also be involved in *Histoplasma* sporulation, as it promotes conidiation in the related thermally dimorphic fungus *Blastomyces dermatitidis* and has been shown to regulate the yeast to hyphal transitions of *Histoplasma* and *B. dermatitidis* [36]. Intriguingly, factors that specifically regulate conidiation without also affecting yeast or hyphal growth have not yet been identified.

In addition to the pathways controlling conidial production, very little is known about the conidia themselves. *Histoplasma* is the only thermal dimorph to produce micro- and macroconidia, but the only known distinguishing criterion is size since nothing is known about molecular composition. Here, we show that both spore types contain chitin, exposed chitin, and mannoproteins. While we observed qualitative differences across strain background and conidiation media, we also observed heterogeneity within the same conidial preparation. It has been shown for other fungi that cell wall heterogeneity within a conidial population can increase the probability in which conidia can be successful in enduring hostile conditions, either in the environment or within host [37]. However, the significance of *Histoplasma* conidial cell wall heterogeneity remains unexplored. In addition to cell wall composition, little is known about gene expression in conidia. We have previously demonstrated that microconidia show distinct transcriptomes from yeast and hyphae [8]. However, it is unknown if the transcriptomes of micro- and macroconidia differ and change during germination. More detailed studies are needed to further characterize these spore types and identify any structural and physiological differences.

Optimization of Histoplasma conidial preparation is foundational for probing this understudied aspect of *Histoplasma* biology. By identifying different conidiation media, it is now possible to investigate the signals that promote conidiation. Finally, by identifying conditions that promote the simultaneous production of micro- and macroconidia, we can now interrogate the differences between these spore types.

## Materials and Methods

### Media and reagents

All media was supplemented with 1x Penicillin/Streptomycin (Gibco). *Histoplasma* yeast were cultured on HMM agarose plates [38] supplemented with 0.5% conditioned media or grown in liquid HMM. Recipes for conidiation media are listed in Supplemental File 1. Conidiation plates were aged for two weeks at room temperature while protected from light, as described for V8 agar [16]. After aging, plates were placed in plastic bags and stored at 4°C until use. Plates were inoculated within two months of pouring. For CFU enumeration, conidia were plated on Brain Heart Infusion (BHI) agar plates (BD) supplemented with 10% sheep’s blood (Colorado Serum Company), 0.05% cysteine HCl (Sigma), 10 µg/mL gentamycin (Thomas Scientific), and 0.5% conditioned media. Plates were incubated at the indicated temperatures for 2 weeks before colony enumeration. Mannose, EDTA, Tween-80, and Triton X-100 were purchased from Sigma Aldrich.

### *Histoplasma* strains and culture conditions

All work with *Histoplasma* yeast was conducted at Biosafety Level 2 (BSL-2), and work with hyphae and conidia was conducted at BSL-3. *Histoplasma* strains G217B and H88 were acquired from American Type Culture Collection (ATCC), and G184AR was a kind gift from Dr. William Goldman (Washington University, St Louis, MO). Strains were inoculated from frozen stocks onto HMM plates and cultured at 37°C, 5% CO_2_ for approximately two weeks. Yeast were subsequently cultured in liquid HMM at 37°C, 5% CO_2_, 150 rpm for one week, with passaging every 2-3 days. For experiments described in Figures 1 and 2, to inoculate conidiation plates, G217B yeast were washed once with D-PBS (Gibco) then diluted in D-PBS to an OD_600_ of 1, and 100 µl was spread per agar plate using glass beads. As G184AR and H88 tend to form large clumps of yeast when grown in liquid culture, we modified our culture preparation for the experiments described in Figure 3. All cultures were spun at 50 x *g* for 20 seconds to pellet larger clumps of cells. The cells remaining in the supernatant were then collected, washed once with D-PBS, resuspended in D-PBS, and sonicated using a Fisher Scientific Sonic Dismembrator Model 100 for 12 1-second bursts. These cells were then diluted in PBS to an OD_600_ of 1, and 100 µl was spread per agar plate using glass beads. After inoculation, conidiation plates were incubated at 25°C for 4 and 6 weeks. The plates were imaged with an Epson Perfection V600 Photo Scanner to show macroscopic hyphal morphology.

### Conidial preparations

Conidia were prepared from six replicate plates and pooled. To prepare conidia, 5 mL of D-PBS was added to each agar plate. The conidia were dislodged by gently rubbing the hyphal surface with a bent glass rod, and the suspension was passed through fiber glass roving (Pyrex) to filter out mycelial fragments. The six duplicate plates were then sequentially washed with 5 mL D-PBS, which was passed through the same glass wool. The conidia were pelleted at approximately 1250 x *g* for 10 minutes and then resuspended in 1 mL or less of D-PBS, depending on pellet size. Conidia were counted on a Axiovert 200 microscope (Zeiss) using disposable Neubauer Improved hemacytometers (INCYTO), and representative micrographs were taken using Micro-Manager [39] version 1.4.21. To assess viability via colony forming units (CFU), spores were serially diluted in PBS containing 0.5% RP-Tamol (Northeast Laboratory) with thorough vortexing, and 500 spores per plate were spread on six replicate BHI plates. Three replicates were incubated at either 37°C or 25°C for 2 weeks before colony enumeration.

### Cell wall staining and microscopy

Cells were fixed in 4% paraformaldehyde (Electron Microscopy Sciences) for 30 minutes then washed twice with PBS. Cells were stained using 50 µg/ml Concanavalin A conjugated to Alexa Fluor 488 (ConA-AF488, Invitrogen C11254) and 100 µg/ml Wheat Germ Agglutinin conjugated to Alexa Fluor 594 (WGA-AF594, Invitrogen W11262) for 1 hour at room temperature on an end-over-end rotator protected from light. Then, calcofluor white was added at the final concentration of 25 µg/ml and incubated for another 10 minutes. Cells were washed once with PBS and resuspended in PBS. Z-stack images were taken on Nikon Eclipse Ti CSU-X1 spinning disk confocal microscope using identical acquisition parameters. FIJI software [40] was utilized to visualize the microscopy data. Representative z-planes are shown in figures. For some images, LUT (look-up table)-adjusted images allowed improved visualization. LUT settings are otherwise constant across images.

### Statistical analysis

Data presented here are from a minimum of 2 replicate experiments. One-way ANOVAs with Tukey’s multiple comparisons tests and simple linear regressions were performed using GraphPad Prism 10. 3-, 4-, and 5-way ANOVAs with post-hoc comparisons using Tukey’s “Honest Significant Difference” method were performed in R. Proportions in Fig 3C were compared using the prop.test function in R.

---

**Figure S1.**
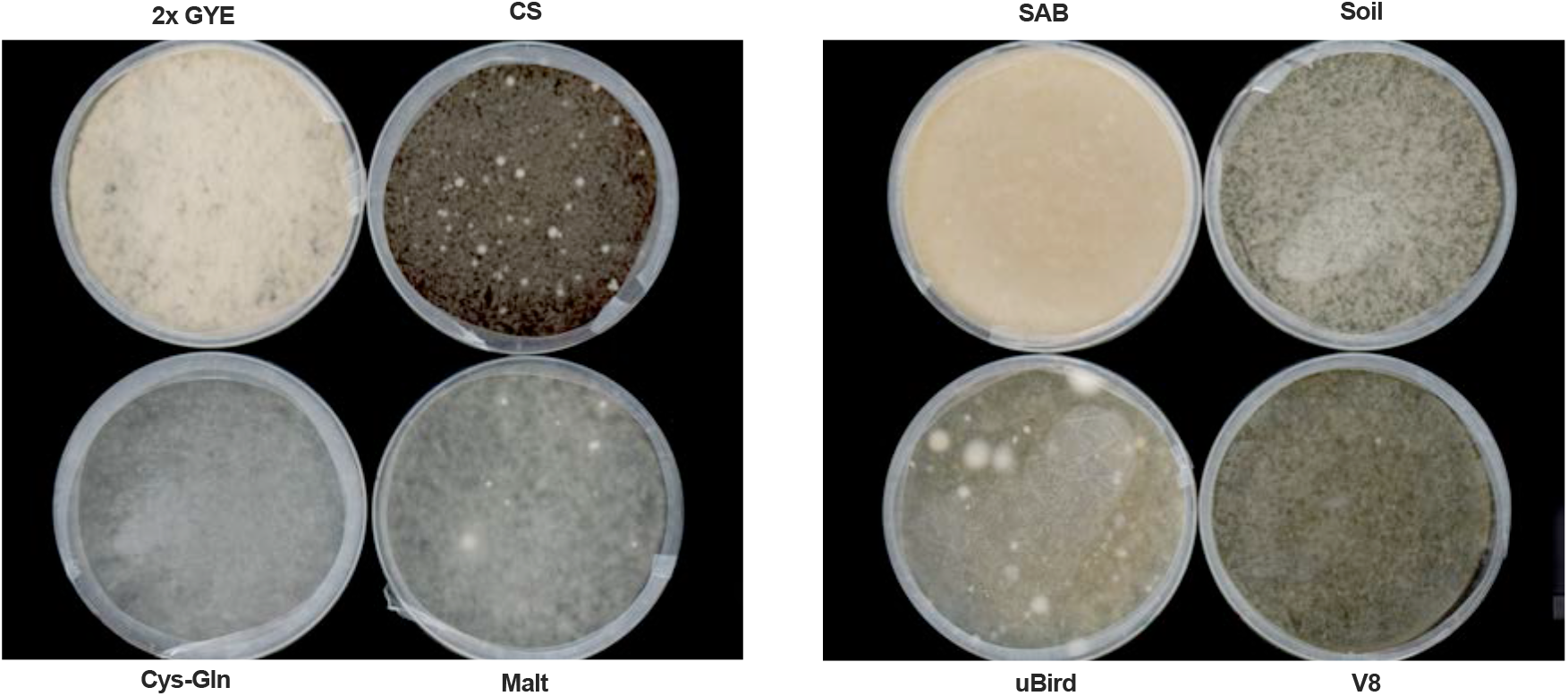
*Histoplasma* grown on different conidiation media has distinct macroscopic appearances. *Histoplasma* yeast were inoculated on agar plates of the indicated media, and images of the agar plates were taken after 4 weeks at 25°C.

**Figure S2.**
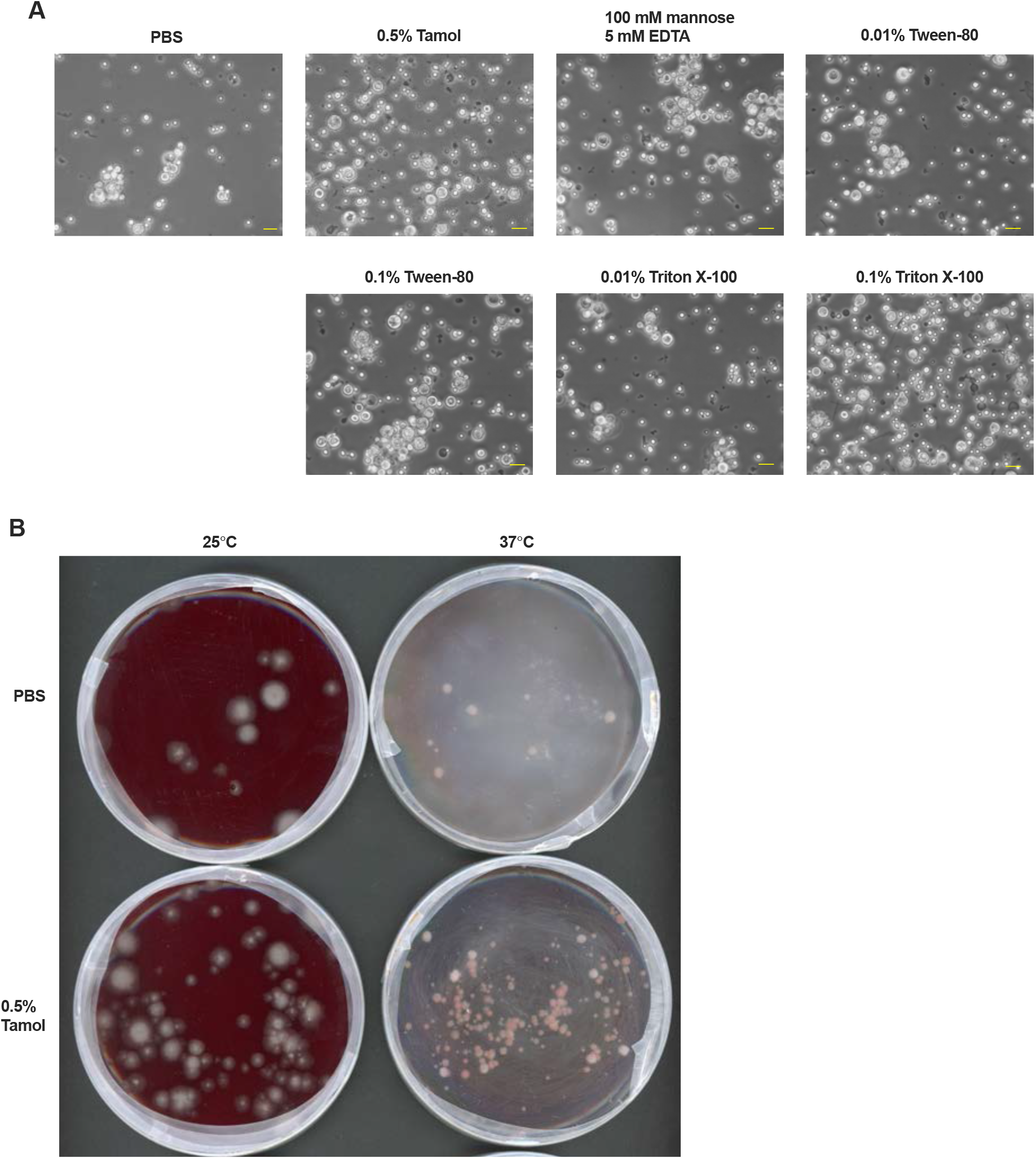
0.5% Tamol disperses conidial clumps and increases plating efficiency. (A) Representative images of conidia generated on V8 agar and resuspended in PBS or PBS with 0.5% Tamol. Yellow scale bar, 10 μm. (B) Conidia generated on V8 agar plates were serially diluted in PBS or PBS with 0.5% Tamol, plated on BHI agar at 500 conidia per plate, and incubated at the indicated temperatures for 2 weeks. Representative images of plates showing increased plating efficiency after diluting in PBS containing 0.5% Tamol.

**Fig S3.**
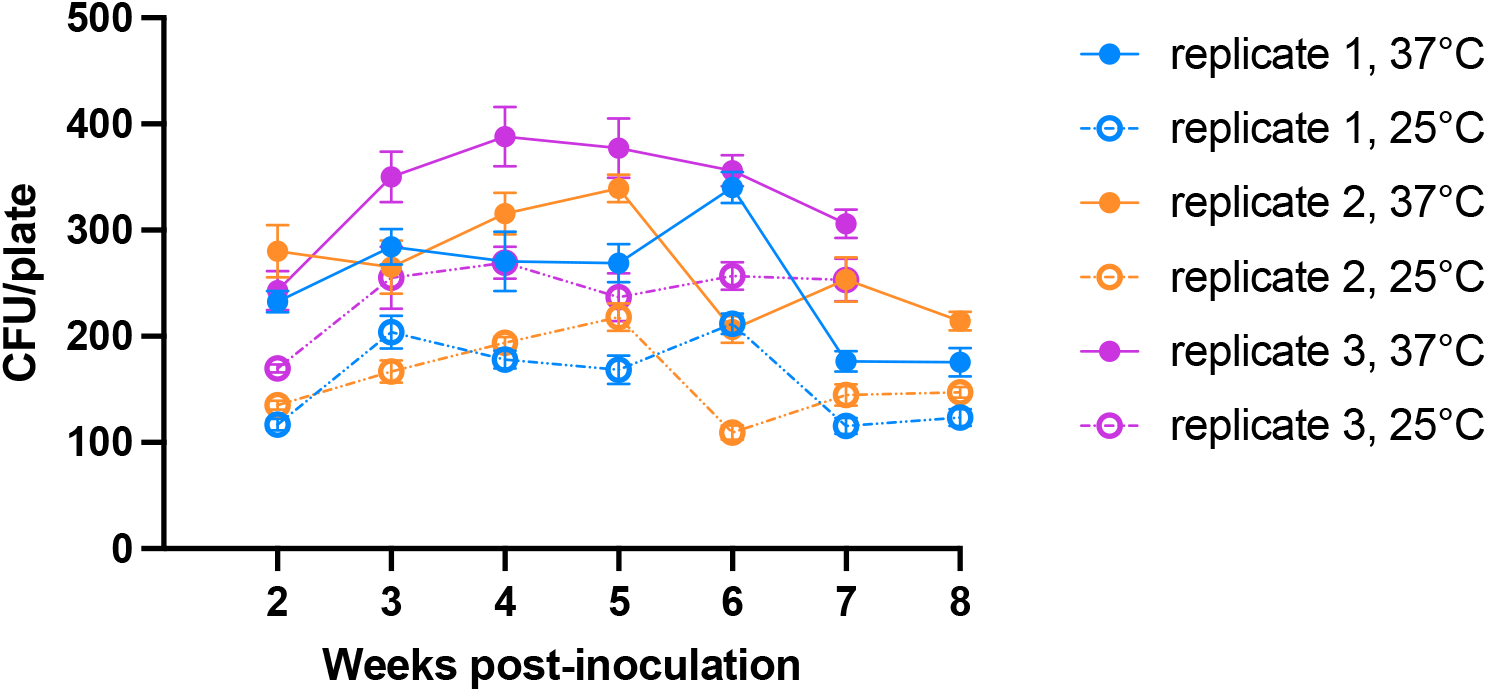
*Histoplasma* conidiation germination rates vary between experiments. Individual replicate experiment data from Fig 2D is shown. Replicates are shown by different colors; solid lines and filled circles are CFUs from spores germinated at 37°C, and dashed lines with open symbols are CFUs from spores germinated at 25°C.

**Fig S4.**
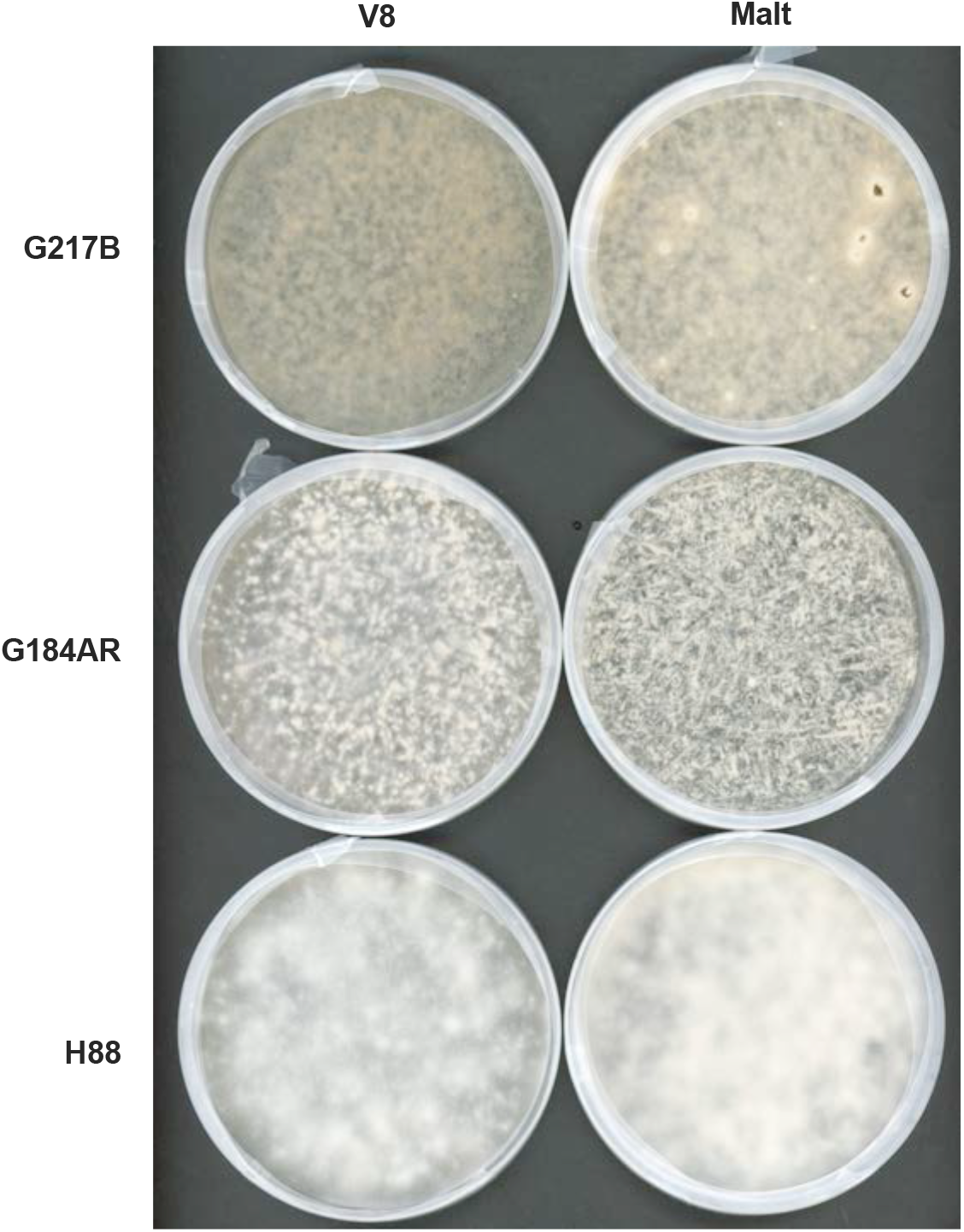
Macroscopic conidiation plate morphology varies with *Histoplasma* strain and media. Yeast from strains G217B, G184AR, and H88 were inoculated on V8 and malt agar plates, and images of the agar plates were taken after 4 weeks of incubation at 25°C.

**Figure S5.**
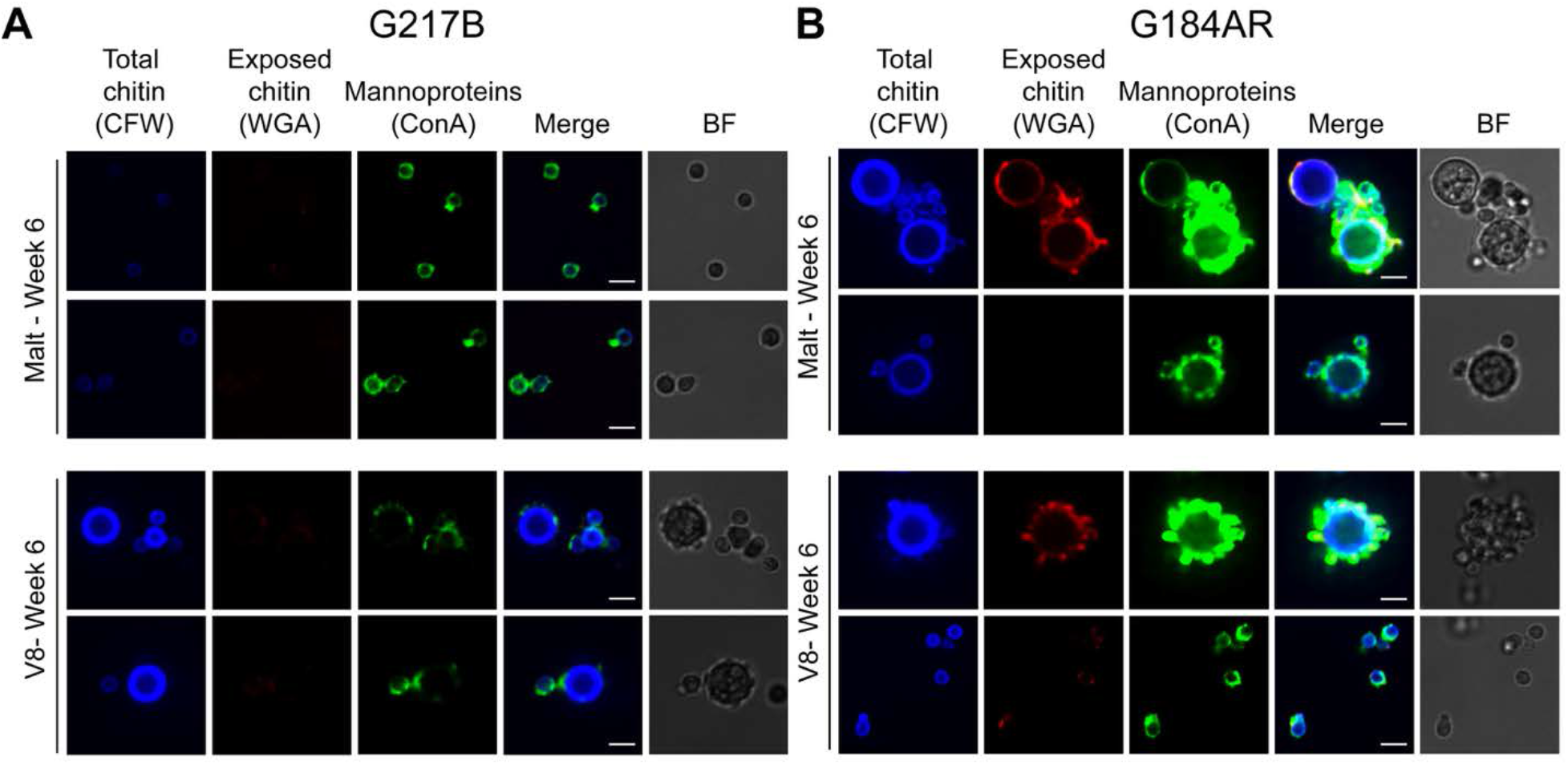
Cell wall composition of conidia at 6 weeks is similar to 4-week samples. The staining panel from Fig 4 was used on conidia prepared after 6 weeks of incubation at 25°C. Representative images for (A) G217B and (B) G184AR conidia are shown. BF, bright field. White scale bars, 5 µm.

**Supplemental Table 1.** Data from Fig 1D were analyzed with a 3-way ANOVA with independent factors for media, temperature, and batch as well as an interaction term between media and temperature; significance was determined by F-test and post-hoc comparisons were made using the Tukey’s Honest Significant Difference method.

|  | Df | Sum Sq | Mean Sq | F value | Pr(>F) |
| --- | --- | --- | --- | --- | --- |
| **media** | 2 | 3.1008 | 1.5504 | 133.04 | < 2e-16 *** |
| **temp** | 1 | 0 | 0 | 0 | 0.994 |
| **batch** | 2 | 0.3744 | 0.1872 | 16.06 | 5.12e-06 *** |
| **media:temp** | 2 | 0.3012 | 0.1506 | 12.92 | 3.52e-05 *** |
| **Residuals** | 46 | 0.5361 | 0.0117 |  |  |

**Supplemental Table 2.** Data from Fig 3B were analyzed with a 4-way ANOVA of log(counts) with independent factors for media, strain, time, and batch. The media and time terms were significant by F-test, and post-hoc comparisons were made using the Tukey Honest Significant Difference method.

|  | **Df** | **Sum Sq** | **Mean Sq** | **F value** | **Pr(>F)** |
| --- | --- | --- | --- | --- | --- |
| **media** | 1 | 0.2239 | 0.22392 | 6.754 | 0.0247 * |
| **strain** | 1 | 0.0514 | 0.05138 | 1.55 | 0.239 |
| **time** | 1 | 0.2923 | 0.29231 | 8.817 | 0.0128 * |
| **batch** | 1 | 0.0067 | 0.0067 | 0.202 | 0.6618 |
| **Residuals** | 11 | 0.3647 | 0.03315 |  |  |

**Supplemental Table 3.**
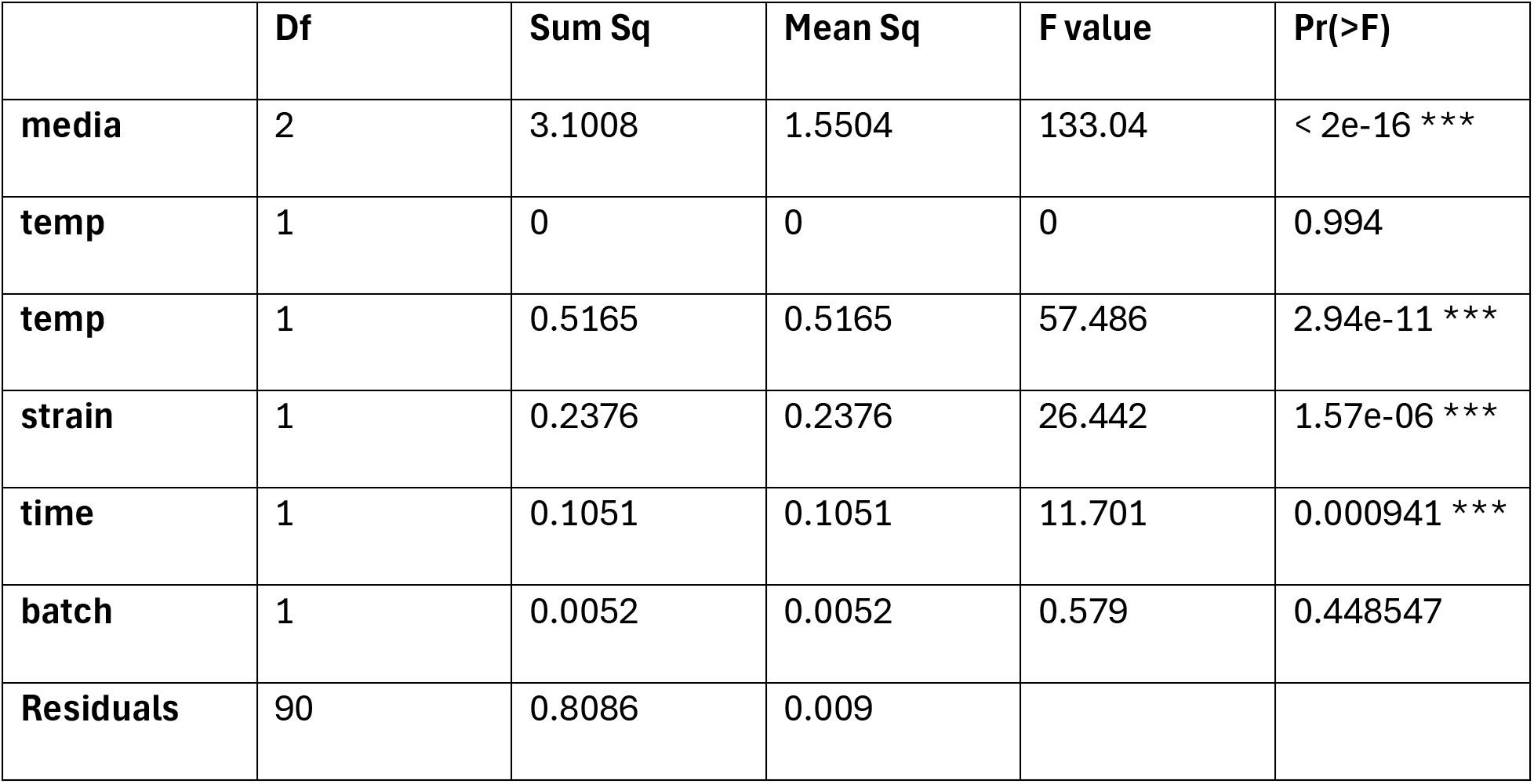
Data from Fig 3D were analyzed with a 5-way ANOVA of log(counts) with independent factors for media, temperature, strain, time, and batch. The media, temperature, strain, and time terms were significant by F-test, and post-hoc comparisons were made using the Tukey Honest Significant Difference method.

## Acknowledgements

We thank Dr. Anna Morrison for sharing the initial conidial preparation protocol. We are grateful to Dr. William Goldman for sharing the *Histoplasma* strain G184AR. We thank Dr. Sinem Beyhan and members of the Sil and Noble labs for helpful discussions.

This research was supported by NIH R37AI066224 (to AS). AS is a Biohub, San Francisco investigator. The funders had no role in study design, data collection and interpretation, or the decision to submit the work for publication.

